# Transcriptomic profile of *Anopheles gambiae* Kisumu mosquitoes infected by neglected malaria parasite *Plasmodium ovale* from gametocyte-carriers in Cameroon

**DOI:** 10.64898/2026.01.27.701957

**Authors:** Daniel Nguiffo‑Nguete, Arnaud Tepa, Aurelie P. Yougang, Francis Nongley Nkemngo, Cyrille Ndo, Stravensky T. Boussougou‑Sambe, Francine Ntoumi, Ayola A. Adegnika, Steffen Borrmann, Charles S. Wondji

**Affiliations:** Parasitology and Microbiology Department, Centre for Research in Infectious Diseases (CRID), Yaounde, Cameroon; Centre for Infection Biology and Tropical Health, Forzi Institute, Limbe, Cameroon; Texas Biomedical Research Institute, San Antonio, Texas, USA; Department of Biological Sciences, Faculty of Medicine and Pharmaceutical Sciences, University of Douala, Cameroon; Institute of Genomics and Global Health, Ede, Nigeria; Fondation Congolaise Pour La Recherche Medicale, Brazzaville, Republic of the Congo; Institut Fur Tropenmedizin, Eberhard Karls Universitat, Tubingen, Germany; German Center for Infection Research, Tubingen, Germany; Centre de Recherches Medicales de Lambarene, Lambarene, Gabon; International Institute of Tropical Agriculture (IITA), Yaounde, Cameroon.; Vector Biology Department, Liverpool School of Tropical Medicine, Liverpool, UK

**Keywords:** *Plasmodium ovale*, *Anopheles gambiae*, transcriptomic profile, upregulated, downregulated, transmission-blocking strategies

## Abstract

Successful transmission of malaria depends on the complex interactions between the Anopheles mosquito vector and the *Plasmodium* parasites. *Plasmodium ovale*, a neglected malaria parasite, successfully develops from ookinete to sporozoite within the *Anopheles* vector. To elucidate the molecular mechanisms underlying this interaction, we compared RNA-seq-based gene expression profiles of *Anopheles gambiae* infected with *P. ovale* and uninfected mosquitoes at 24 hours, 9 days, and 17 days post-infection. The results showed that 2,885 *P. ovale* transcripts were present only 24 hours after infection. During ookinete invasion (24 h post-infection), differential gene expression analyses revealed the up-regulation of genes related to metabolic processes and the down-regulation of genes associated with cytoskeletal activity in the mosquito. Notably, the non-immune genes with unspecific function AGAP003776, (Fold Change, FC 132.0), AGAP003777, (FC 88.3), and AGAP003778, (FC 104.1), Troponin C (Fold Change, FC 85) and Myofilin (FC 33.3) exhibited the most significant overexpression. Among the immune genes that were upregulated CTL3 (FC 55.9), CLIPB12 (FC 49.4), CTLMA5 (FC 14.5), TRYP7 (FC 24.4), CLIP C9 (FC 12.1) TRYP5 (FC 12.2), LRIM10 (FC 11.2), PPO6 (FC 7.7). This initial analysis of the interaction between *P. ovale* and *An. gambiae* identified several well-known candidates for transmission-blocking strategies, including LRIM1, APN1, and D7 family proteins. In addition, new potential candidates, including AGAP003776, AGAP003777, and AGAP003778 cluster, CLIPB12, LRIM10, the APN cluster, AGAP004860, ABCC9, CYP9K1 and GSTD3 were identified. These potential new candidate genes could play a significant role in the development of transmission-blocking strategies for *An. gambiae* infected with *Plasmodium*, particularly *P. ovale*. The urgent functional validation of these genes is required.

## Introduction

Malaria remains one of the most severe infectious diseases in the world. Most effort to control and eliminate malaria mainly focus on elimination of *Plasmodium falciparum* parasites which is the dominant species in Sub-Saharan Africa and responsible for highest morbidity and mortality (1, 2). Between 2000 and 2015, global malaria incidence and mortality declined by 37% and 60% respectively. This decline was largely due to the widespread deployment of effective control interventions including long-lasting insecticidal nets (LLINs), artemisinin-based combination therapies (ACTs), seasonal malaria chemoprevention (SMC) (2, 3). These methods primarily target *P. falciparum* parasite. However, recent molecular surveillance data is revealing the increase and persistent circulation of non falciparum malaria parasites in Africa particularly *P. malariae, P; vivax* and *P. ovale* (4-6). Despite significant reduction in the disease burden, this progress has stalled since 2015, challenged by widespread emergence of insecticide and antimalarial drug resistance, diagnostic failure from pfhrp2/3 deletions, as well as the emergence of invasive mosquito species, jeopardizing future progress toward malaria elimination. Together, these factors highlight the urgent need for complementary strategies to accelerate malaria elimination (2, 7, 8). These additional interventions could include transmission-blocking approaches that target parasite development in the mosquitoes, and genetic replacement of vector populations with non-vectors populations with the aim to interrupt parasite transmission by creating mosquitoes resistant to *Plasmodium* infection through gene drives (9, 10).

Enhanced molecular surveillance has demonstrated that non-falciparum species, particularly *P. malariae* and *P. ovale* play a significant role in sustaining malaria reservoir in Africa. Coinfection rates with these species can reach up to 50% in certain regions (11). Furthermore, persistent transmission of *P. malariae* and *P. ovale* has been observed in area *P. falciparum* transmisision is declining (12, 13). Recent molecular surveillance in Cameroon, has recently confirmed that the non-falciparum species such as *P. ovale curtisi, P. ovale wallikeri*, and *P*.*malariae* are circulating, either in mixed or mono-infections with *P*.*falciparum*. These infection have been documented accross several regions in both symptomatic and asymptomatic individuals (5, 14, 15). Crucially, *P. ovale* shares with *P. vivax* the ability to form dormant liver stages (hypnozoites), which may reactivate months or years later to produce relapses if not properly treated (16). This biological feature not only prolongs infection but also sustains transmission reservoirs that bypass *P. falciparum*-focused interventions. Actually, little is known about its epidemiology, relapse dynamics, or interactions with mosquito vectors. The lack of continuous culture systems severely limits biological studies, leaving fundamental questions unanswered.

Several studies in *Anopheles gambiae* vector have shown that the mosquito’s immune response against *Plasmodium* infection is complex and triggers the defense factors like Thioester-containing protein 1 (TEP1), gambicin, and Nitric Oxide Synthase (NOS). These factors mediate defense against both *P. falciparum, P. vivax* (human malaria) and *P. berghei* (rodent malaria) (17-19), while other genes like AgMDL1 and FBN39 show specificity for *P. falciparum* resistance (17). In addition, the RNA-interference based gene silencing (RNAi) of TEP1, LRIM1, APN1 showed the variation in response which can increase or decrease mosquito susceptibility to infection, implicating them as transmission-blocking factors target for *P. falciparum, P. vivax* and *P. berghei* (17, 20, 21). Whether similar mechanisms act against *P. ovale* is unknown. Recent transcriptomic analyses of *P. ovale* have provided initial insights into parasite gene expression during the blood stage (22). However,there remains limited understanding of mosquito immune responses or transmission-blocking factors specific to *P. ovale*.

To start filling this gap, this study experimentally infected *An. gambiae* Kisumu laboratory mosquitoes colony with field collected *P. ovale* and performed a transcriptomics analysis to identify innate immune factors involved in the mosquito’s response to *P. ovale* and putative mechanisms of parasite immune evasion. This study identified the potential candidate genes for transmission-blocking strategies against *P. ovale*, which may complement existing *P. falciparum* tools to support malaria elimination in Africa.

## Material and method

### Ethical statement

Ethical approval was obtained from the National Ethics Committee of Cameroon under the following permit N°2022/08/1480/CE/CNERSH/SP. All human volunteers were enrolled after written informed consent was signed from the participant and/or their legal representative.

### Study area

Parasitological surveys were conducted in several villages in the Centre Region of Cameroon to identify carriers of *P. ovale* gametocytes, following preliminary findings indicating the presence of non-*Plasmodium falciparum* species circulating within the population (23)

### Mosquitoes rearing

*Anopheles gambiae* Kisumu laboratory mosquitoes were reared at the Centre for Research in Infectious Diseases (CRID) under standard conditions of 28 +/-2°C and 85 +/-5% humidity with a 12 h day/night cycle according to standard rearing procedures (24). Adults were kept in paper cups and fed on 10% sucrose.

### Identification of gametocyte carriers

Thick and thin blood smears collected from the finger of each participant enrolled in the study were air-dried and stained with 10 % Giemsa solution to detect of *Plasmodium* parasites at 100X objective. Gametocyte density was quantified microscopically by counting against 500 leukocytes, assuming a standard white blood cell count of 8,000/μl (25).

### Experimental infections

A volume of 4 ml of blood was collected from gametocyte carriers. The blood was then centrifuged at 2000 rpm for 3 minutes at 37°C and the serum was replaced with an equivalent volume of non-immune AB serum (Sigma-Aldrich, Taufkirchen, Germany). Groups of 100 adult female mosquitoes aged between 5 and 7 days, starved 12 hours before the experiments, were placed in paper cups covered with mosquito netting. A volume of 450ul of reconstituted blood was added to a glass feeder covered with a parafilm membrane and maintained at 37°C using a thermostatic water bath (Fisher Scientific INC, Isotemp 4500H5P, Pittsburgh USA). Mosquitoes were fed in the dark for 45 min; thereafter, unfed and partially fed mosquitoes were removed and killed. The fully fed mosquitoes were kept in the insectary at 26 ± 2°C and 70-80% relative humidity, with a constant supply of 10% sucrose soaked on cotton. On day(s) 1 (24 hr), 9 and 17 post-infection, surviving females were stunned by exposure to ice for approximately 1 min. They were then dissected in a drop of 0.4% diluted mercurochrome and the stained midgut was examined on day 9 for the detection and quantification of oocysts under a light microscope. The dissection was not performed on day 17 because, after preservation in RNAlater, too few mosquitoes were left. However, PCR analysis of the head and thorax on 20 samples confirmed the presence of *P. ovale* mono-infection at this different stages.

Mosquitoes from this infection were preserved in RNA later on day 1 (24-28h, during invasion of the intestinal epithelium), day 9 (presence of oocysts) and day 17 (presence of sporozoites) for subsequent transcriptomic studies. The same protocol was used to feed control mosquitoes with blood from a non-infected individual.

### Validation of Plasmodium species after infection by PCR

DNA was extracted from the abdomen of mosquitoes on day 9 (for the presence of oocysts) and from the head and thorax on day 17 (for the presence of sporozoites) using Livak’s protocol (26). Nested PCRs were then performed to validate the presence of *P. ovale* oocysts and sporozoites in the infected samples (27). We did not perform PCR on the fed mosquitoes on day 1 (24 h) post-infection because at this stage the blood of the infected person is still present in the abdomen of the mosquito and does not guarantee the success of the infection.

### RNA extraction and sequencing

Total RNA was extracted at CRID from *An. gambiae* (*Kisumu* laboratory strain) infected with *P. ovale* using the PicoPure RNA Isolation Kit (Life Technologies, Carlsbad, CA, USA) and stored at -80°C. For each time point, three groups of 10 mosquitoes were used (infected and uninfected). Quality and quantity of RNA obtained were assessed using a “NanoDrop Lite” spectrophotometer (Thermo Scientific Inc., Wilmington, USA) and a Tapestation system (Agilent Technologies, Inc., Santa Clara, United States).

Library preparation, sequencing and initial data quality control were carried out by Novogene, UK. Methods used for ribosomal RNA-depletion, library preparation, and sequencing were performed as previously described (28). Samples were sequenced using the NovaSeq™ 6000 Sequencing System (Illumina, Inc., San Diego, CA, USA) by the Novogene Company (Cambridge, UK).

### RNA-seq data analysis

The analysis began by performing quality checks on each FASTQ file using the FastQC tool (Andrews, 2010), followed by aggregating the results into a comprehensive report with MultiQC (Ewels et al., 2016). The gene expression profile was analyzed using the DESeq2 R package (29). The genes were counted as differentially expressed if FDR-corrected P-values *<* 0.05 and absolute fold-change values >1 when comparing *P. ovale* infected to uninfected.

Plots were generated with the log (FC) value using R software to visualise the expression level. The *p-values* used to evaluate significantly enriched GO terms were calculated based on Fisher’s exact test and corrected by the Benjamini-Hochberg multiple test correction method. Finally, we used a FDR adjusted *p*-value *<*0.05 to tag statistically significant overrepresented GO terms associated with the list of differentially expressed genes (DEG).

To identify potential candidate genes that can be linked to reduce *Plasmodium* transmission, we used several approaches including the significant upregulation in response to infection (17), high expression levels (Read Counts)(30), relevant functional annotation (Gene Ontology) (31) and functional evidence from literature (32).

### Quantitative reverse transcriptase PCR

RNAseq expression patterns of 13 of the most over-expressed immune genes were validated by quantitative reverse transcription PCR (qRT-PCR), including TEP4 (AGAP010812), CTL4 (AGAP005335), TEP12 (AGAP008654), LRIM10 (AGAP007455), CLIP B12 (AGAP009217), Trypsin7 (AGAP008293), Myofilin (AGAP004161), Troponin C (AGAP003777), TEP1 (AGAP010815), CTLMA2 (AGAP005334), PGRPS2 (AGAP006343), PPO2 (AGAP006258), TOLL1A (AGAP001004). The primers are listed in S1 Table. Briefly, 1mg of total RNA was used for cDNA synthesis using Superscript III (Invitrogen, Carlsbad, CA, USA) with oligo-Dt20 and RNase H according to the manufacturer’s instructions. The qRT-PCR amplification was performed following standard protocol after establishing the standard curves for each gene to assess PCR efficiency and quantitative differences between samples using serial dilution. The relative expression of each gene was calculated according to the 2^_ΔΔCT^ method and compared between the two strains after normalization with the housekeeping genes ribosomal protein S7 (AGAP010592) and *Elongation Factor* (AGAP009441) (33).

### Polymorphism Analysis in Plasmodium ovale infection

RNA-seq variant calling was performed using Samtools mpileup and Varscan 2 (Koboldt et al., 2013) as previously described in the EasyRNAseq Repository (34). Variants were functionally annotated with Snpeff (35). Diversity metrics (Theta π, Theta Watterson, Tajima’s D) and FST were computed from sync files with Grenedalf, using per-sample and total minimum read depths of 30. Nonsynonymous SNP associations between infected (Inf) and uninfected (Uninf) pools were evaluated with Fisher’s exact test (PoPoolation2), requiring min-count ≥ 5 and min-coverage ≥ 10 per pool. The analysis of sample clustering according to their phenotype was performed using Plink 1.9.

## Results

### Parasitological survey

For this work, 293 individuals were sampled for gametocyte detection and experimental infection. All the individuals were asymptomatic (body temperature <37.5°C). The overall prevalence of *Plasmodium* infection was 55% (160) with the higher prevalence obtained with *P. falciparum* 50% (145), followed by coinfection with *P. malariae/P. ovale* 14% (41). The prevalence of *P. ovale* gametocyte carriers monoinfection identified by blood smear was 0.3% (1).

High infection rates after six experimental infections of *An. gambiae* with *P. ovale* were carried out in the field using *P. ovale*-infected blood from different gametocyte carriers. In total, 1560 (91%) *An. gambiae* Kisumu were fully fed with infected blood through an artificial parafilm membrane. *Plasmodium ovale* gametocytes were infective for two experiments (one with *P. ovale* mono-infection (S1 Fig) and the second with *P. falciparum* and *P. ovale* co-infection). This study only considered cases of *P. ovale* mono-infection. Prevalence of Kisumu mono infection was 100% (31 infected out of 31 mosquitoes dissected) with total oocyst counted in infected midguts was 846 with the number of oocysts observed in a midgut ranged from 3 to 51 oocysts.

### Differential transcriptional expressed genes in *An. gambiae* Kisumu strain

Quality control metrics revealed an average sequencing depth of approximately 60 million reads per sample, with GC content consistently around 50% and duplication rates ranging from 74% to 96%. Mapping success varied across samples, with most achieving alignment rates above 80%, though a few outliers such as InfD9_1_R1, showed markedly lower alignment efficiency (S2 Table). The PCA plot illustrated the gene expression profiles of *An. gambiae* mosquitoes at different time points post-infection (Fig 1a). On day 1 post-infection, both infected (D1. Inf) and uninfected (D1.Uinf) mosquitoes formed distinct clusters, indicating significant differences in gene expression shortly after infection. By day 9, the infected (D9.Inf) and uninfected (D9.Uinf) mosquitoes showed similar clustering, suggesting some overlap in gene expression profiles, yet still maintaining clear distinctions with the infected and uninfected day 1. On day 17, the infected (D17.Inf) and uninfected (D17.Uinf) mosquitoes exhibited a similar clustering pattern to day 9.

**Fig 1:**
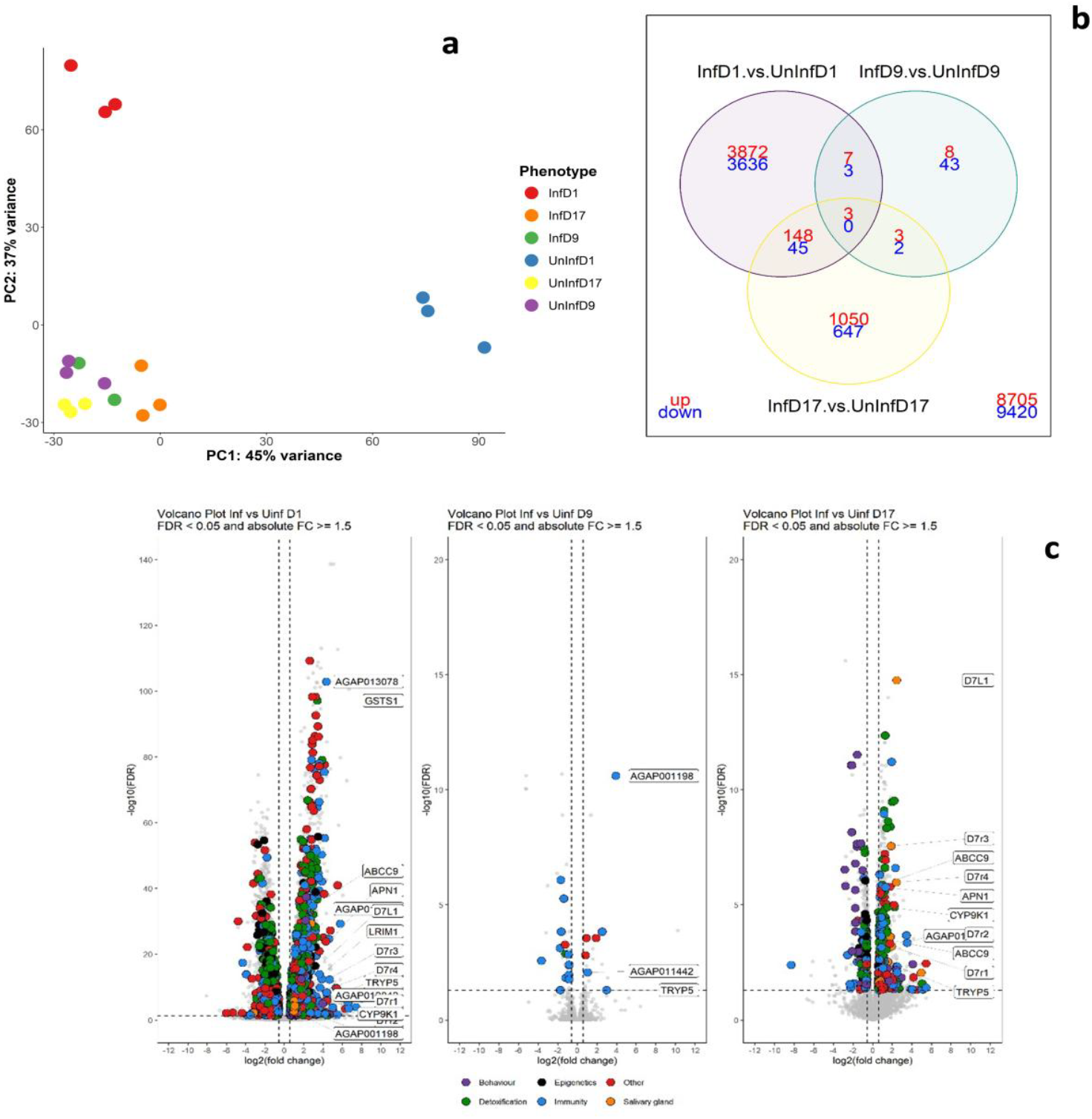
**a**-Principal component analysis (PCA) of transcriptome variation in *Kisumu* mosquitoes at three time point (day 1, day 9 an day 17 post feeding). **b**-Venn diagram showing the repartition of genes expression at different time points. **c**-Volcano plots showing *An. gambiae* gene expression at three different time points after feeding on *Plasmodium ovale*-infected blood versus uninfected blood.

### Gene differentially-expressed in Kisumu according to different time points

During the initial *P. ovale* invasion, 24h after feeding with infected blood, 4030 genes were over-expressed in infected mosquitoes compared to control (blood fed with uninfected blood) and 3684 were down-expressed (Fig 1b). Analysis of the list of transcripts upregulated revealed that the AGAP006622 was the top highest overexpressed with 235.5 FC. Interestingly, three genes with consecutive numbers, no putative function and predicted gene ontology (AGAP003776, FC (132.0), AGAP003777, FC (88.3), and AGAP003778, FC (104.1) were among the top upregulated genes with consistant read count (S3 Table). Among the annotated genes, Troponin C gene (AGAP006181) has the highest FC (85.1) and is a calcium-binding protein that plays an important role in the regulation of muscle contraction by binding calcium ions (Ca^2+^) (S3 Table). Besides, another non-immune gene Myofilin (FC=33.3) implicated in mosquito flight was significant overexpressed. Out of 50 top significant expressed genes, thirty five had unknown functions. The most immunes/digestive enzymes differentiate expressed genes with high expression level were CTL3 (FC 55.9), CLIPB12 (FC 49.4), CTLMA5 (FC 14.5), TRYP7 (FC 24.4), CLIP C9 (FC 12.1) TRYP5 (FC 12.2), LRIM10 (FC 11.2), PPO6 (FC 7.7) (Fig 1c). Furthermore, some overexpressed immune genes were significantly down expressed either at Day 9 (PGRPS3, CLIPB15, LRIM1, TEP1) or at Day17 (AGAP010774, CLIPB3, CLIPB7, LYSC4,) (S3 Table).

Among the down-regulated genes at 1dpi, twelve were associated to immune response including AGAP006722 (FC -5.7) and CLIPD8 (-5.0). In addition, CTLSE2 and TEP3 were up-regulated at 17dpi.

The results also showed a significant downregulation of key genes within the Toll-like receptor pathway at 1dpi including REL1 (FC -1.5) belonging to Toll pathway and REL2 (FC -3.0) from immunodeficiency (IMD) pathway, both of which are critical regulators of mosquito immunity against *Plasmodium* infection. Combined suppression of REL1 and REL2 in *An. gambiae* infected with *P. ovale* resulted in low expression of LRIM1 and TEP1 genes, two essential components of the complement-like immune complex which was subsequently suppressed at 9 dpi. This sustained downregulation correlates with a compromised immune defenses, contributing to the high *P. ovale* infection rate (98%) observed in this study. These findings underscore the essential roles of REL1 and REL2 pathways in modulating effective antiparasitic immunity and highlight their potential as targets for interrupting malaria transmission.

During oocyst development and maturation on day 9 post infection, before differentiation and release of sporozoites in mosquito haemocel, 21 genes had higher expression and 48 genes had lower expression levels in infected mosquitoes (Fig 1b). Beside, immune gene (HPX3, FC=3.7), digestive gene (Trypsin 5, FC=8) and vitellogenin (Vg) (FC=3.0) was among the overexpressed genes. Among the down-regulated genes at 9dpi, AGAP005310, FC =-12.6, was the most significant but was upregulated at 1dpi and annotated as a Chymotrypsin.

At 17dpi during the salivary gland invasion by sporozoites, 1204 genes were upregulated and 694 were downregulated. The top overexpressed genes belong to the salivary protein (Fig 1c). Most of the genes up-regulated at 17dpi were down regulated at 1dpi (689/840(82.0%). Two immune genes TEP13 and CTLSE2 were up-regulated at 17dpi and down-regulated at 1dpi and 9dpi. The list of down regulated genes is found in file 5.

### Identification of transmission blocking strategy within the genes commonly upregulated in different groups

#### Gene expression profiles across infection time points

To identify candidate genes potentially linked to transmission blocking strategy, we analysed gene expression profiles both within and between the different infection time points and then associated with gene ontology (GO) terms.

Among the immune system-related genes found at day 24 hours post-infection, several key immune genes showed significant upregulation. CTL3 (FC: 55.9) and CLIPB12 (FC: 49.4) demonstrated remarkable overexpression with extremely high read counts (119,505.1) in CLIPB12 infected group, followed by other immune effectors including CLIPC9 (FC: 12.1), LRIM10 (FC: 11.2), PPO6 (FC: 7.7), TEP12 (FC: 5.3), TEP1 (FC: 3.8). Alongside, Leucine-Rich Repeat Immune Protein 1 (LRIM1) gene was significantly upregulated 24 hours after infection (FC: 7.0), consistent with its established role in blocking malaria invasion. LRIM1 had high read count (23,327) further confirming its importance during early infection. In addition, a cluster of aminopeptidase N (APN) genes showed significant upregulation, including APN1 (FC: 3.8), APN2 (FC: 8.3), APN3 (FC: 4.8), APN4 (FC: 6.2), and APN5 (FC: 8.5). APN1 have being well established as transmission-blocking vaccine target.

#### Genes Upregulated at Day 1 and Day 9 (Early and Mid-stage Infection)

Analysis of genes upregulated at both Day 1 and Day 9 identified seven genes commonly upregulated at both time points. Two proteolytic enzymes directly linked to proteolysis and immune response: AGAP011442 (Serine carboxypeptidase 1) showed significant upregulation at both Day 1 (FC: 5.1) and Day 9 (FC: 2.0) with substantial read counts (8,245 at Day 1) (S4 Table). Moreover, AGAP001198 (Chymotrypsin) exhibited moderate upregulation at Day 1 (FC: 2.4) and strong upregulation at Day 9 (FC: 15.1), suggesting a potentially important role during mid-stage.

#### Genes Upregulated at Day 1 and Day 17 (Early and Late Infection)

Comparison of gene expression between Day 1 and Day 17 revealed 148 common genes (S4 Table). Known transmission-blocking candidates including APN1, and the D7 family proteins (D7r2, D7r3, D7r4, D7L1, D7r1) were consistently upregulated at both time points. Moreover several novel candidates showed promising expression patterns, including: AGAP004860 (M1 zinc metalloprotease) maintained strong upregulation at both Day 1 (FC: 11.5) and Day 17 (FC: 2.4) with exceptionally high read counts (53,103 at Day 1). ABCC9 (ATP-binding cassette transporter) showed consistent upregulation at Day 1 (FC: 4.5) and Day 17 (FC: 2.0) with substantial read counts (more than 1000 in infected group). At the end, CYP9K1 (Cytochrome P450) displayed upregulation at both Day 1 (FC: 2.3) and Day 17 (FC: 3.8) with significant read counts upper than 1000 in infected group. Moreover, four of these genes were down-regulated at 9dpi (S4 Table).

#### Genes upregulated across all time points

Three genes were found upregulated at all three time points (Day 1, Day 9, and Day 17). Among these three genes, Trypsin 5 (AGAP008291) with FC of 12.2, 7.8 and 33.6 respectively at Day 1, Day 9, and Day 17 maintained consistent upregulation, though with moderate read counts while the AGAP012842 (unspecified product) displayed upregulation across all time points, with low read counts (S3 Table).

#### Gene ontology enrichment

GO term enrichment was analyzed at 24 hours, day 9 and day 17 days after the infection with infected blood meal (Fig 2).

**Fig 2.**
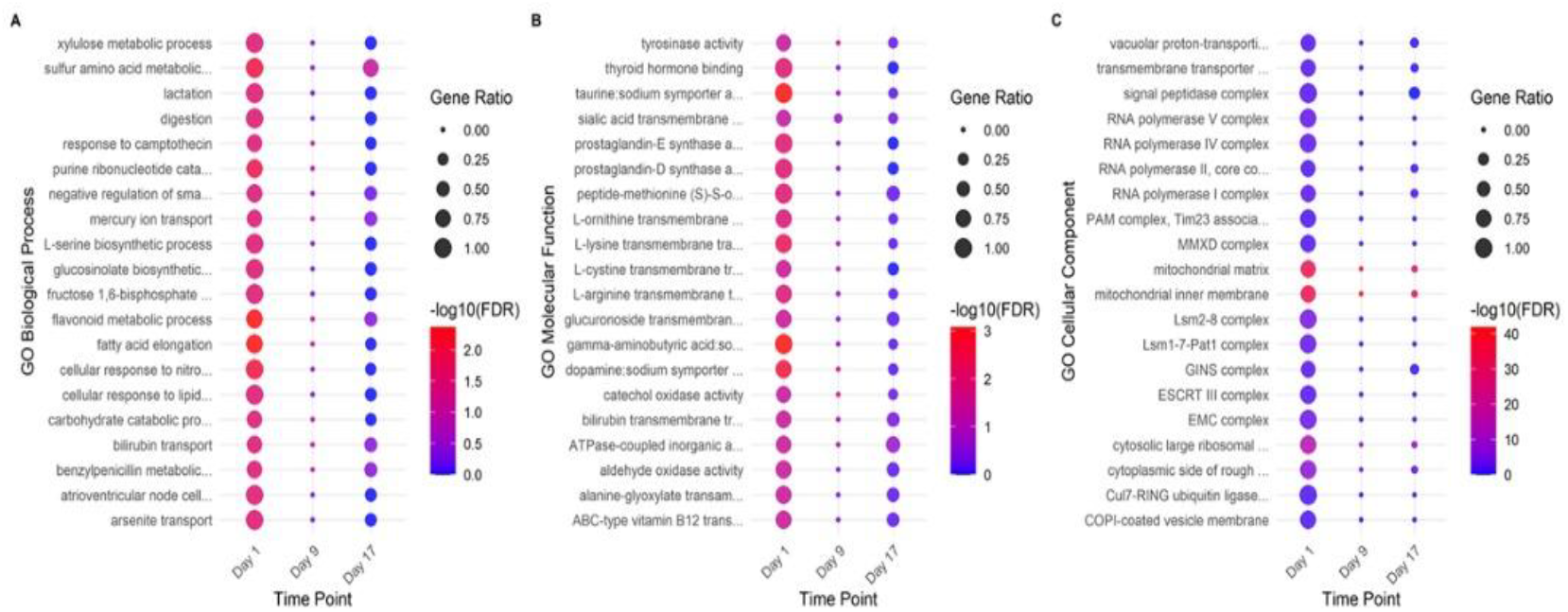
Enriched Gene Ontology terms among differentially expressed genes in infected mosquitoes compared to uninfected ones. A) GO Biological process, B) GO Molecular function, C) GO Cellular component

At 24 hours, biological process revealed that *metabolic process* was the most GO term for up regulated genes in *P. ovale* infected mosquitoes, followed by translation, while *cytoskeleton* was the most down regulated GO term, followed by *endocytosis* and *cell-cell adhesion*. Among the most significant genes, with high expression rates in infected mosquitoes, we had the mitochondrial function. Enriched molecular function of up regulated genes included structural constituent of ribosome, and electron transfert activity while for down regulated genes the *protein kinase activity* was predominant (Fig 2).

At day 9, the most altered GO term for up regulated genes in *P. ovale*-infected mosquitoes was *positive regulation*, and the most altered GO term for down regulated genes was also *defense response to other organism*.

At day 17 day post infection (dpi), biological process revealed that *multicellular organismal process* was the most GO term for up regulated genes in *P. ovale* infected mosquitoes, followed by *G protein-coupled receptor* signaling pathway, while *response to stimulus and sensory perception of smell* was the most down regulated GO term. Among the most significant genes, with high expression rates in infected mosquitoes, we had the cellular response to oxygen-containing compound. Enriched molecular function of up regulated genes included structural constituent of *G protein-coupled receptor* activity while for down regulated genes the *odorant binding* was predominant (Fig 2).

#### Identifying mosquito genes that can block transmission using the Gene Ontology approach

Transcriptional analysis of *An. gambiae* infected with *P. ovale* revealed significant gene upregulation at different time points post-infection. Through analysis of Gene Ontology (GO) terms and expression patterns, we identified several promising candidates genes for transmission-blocking strategies.

The relationship between the gene with high expression and relevant GO terms, have been established and the results showed that, ANP1 and LRIM1 were all upregulated during the infection and belong to various GO and were well established to limiting parasites invasion. The gene like CLIPB12 was found sharing the same GO, Proteolysis (GO:0006508), Immune response (GO:0006955), with LRIM1 and APN1. LRIM10 gene upregulated at D1 was found sharing the same GO with LRIM1, Immune response (GO:0006955), Extracellular region (GO:0005576) and Protein binding (GO:0005515) (S4 Table). AGAP004860 upregulated at 1dpi and 17dpi sharing with APN1, ClipB12 the same GO: Metallopeptidase activity (GO:0008237) and Proteolysis (GO:0006508) (S5 Table). The complex of APN were all upregulated and shared the same function with APN1. ABCC9 gene was found belonging to Transporter activity (GO:0005215). Other genes (CYP9K1, GSTD3) implicated in detoxification and oxidative stress response were also upregulated at Day1 and D17 belonging to the Oxidoreductase activity (GO:0016491) (S5 Table). The gene AGAP001198 upregulated at Day1 and Day9 is associated with GO terms of Proteolysis (GO:0006508) and Serine-type endopeptidase activity (GO:0004252) directly shared with CLIPB12 and APN1 (S5 Table). APN2, APN3, APN4, and APN5 were all upregulated and sharing the same GO with APN1 (S5 Table). A cluster of uncharacterized genes (AGAP003776-AGAP003778) shows exceptionally high fold changes (FC 88.6 - 132). Although they lack precise GO annotations, their strong co-expression with immune and proteolytic genes strongly suggests they may play a role in host-parasite interaction or immune modulation. AGAP003773 to AGAP003778 was found up-regulated during early infection in mousquitoes infected by O’nyony-nyong Virus (Joanna Waldock, 2010). Their marked induction across infection stages warrants further functional characterization as a novel transmission-blocking candidates.

#### qRT-PCR validation of relative expression levels estimated by RNA-Seq

The qRT-PCR was performed to validate the FC of thirteen genes, namely TEP4, CTL4, TEP 12, LRIM10, CLIP B12, REL2, PGRPS2, PPO6, TOLL1A, CTLMA2, Trypsin7, Myofilin, and Troponin C (Fig 3). Their qPCR expression was similar to RNAseq profile, with either high or low expression at 24 hours post infection.

**Fig 3:**
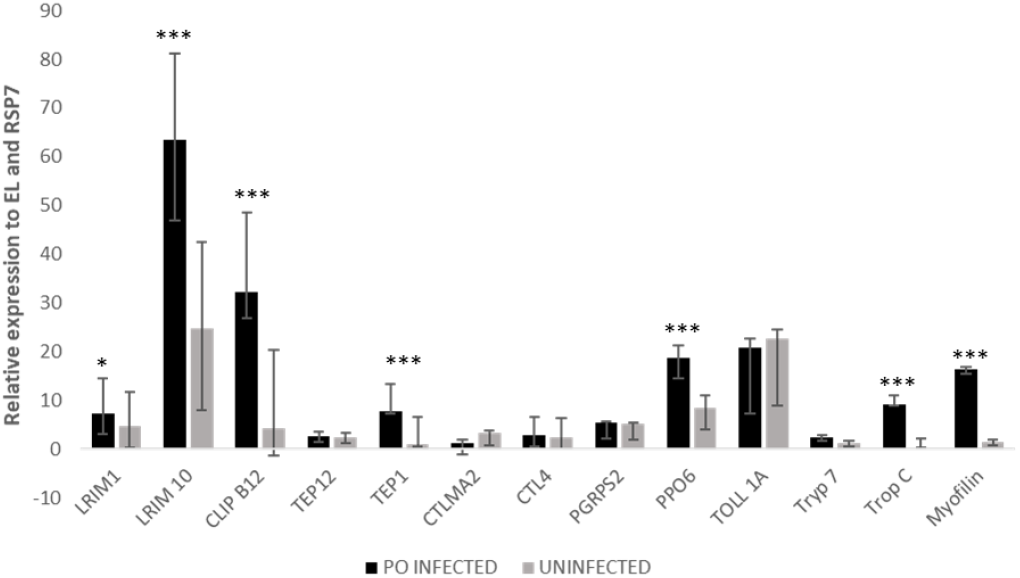
Relative transcript levels of immune genes in adult female *Anopheles* Kisumu at 24H post infection.

#### Early selection on immune genes following *Plasmodium ovale* infection

To investigate genetic polymorphisms linked to mosquito susceptibility to *P*.*ovale* infection, variant calling was performed across multiple post-infection time points (S2 Fig) and genes with FST ≥ 0.1 between infected and uninfected pools at Day 1 were prioritized, identifying immune-related candidates such as LRIM17, CLIPA9, CLIPB1, CLIPC7, REL1, and CLIPB3 (Fig 4). Differentiation was strongest for CLIPB3 (FST = 0.46 at Day 1, 0.51 at Day 9) and CLIPA9 (0.39), indicating marked divergence in allele frequencies. Tajima’s D and nucleotide diversity patterns complement these findings: negative Tajima’s D in infected pools for CLIPB1, CLIPB19, and REL1 (up to −4.01) suggests directional selection, while positive values for LYSC7 and CLIPB3 at later time points indicate balancing forces. Diversity was generally lower in infected pools for CLIPB genes, contrasting with higher diversity in CLIPA9, consistent with locus-specific evolutionary pressures. The heatmap (Fig 4) highlights nonsynonymous SNPs within genes under selection, including LRIM17 (Pro401Ala, Tyr234His), CLIPA9 (Gln25Lys, Ile179Met, Phe233Tyr), CLIPB1 (Leu22Ser), and CLIPC7 (Ser471Thr), which correspond to loci with elevated FST and Tajima’s D shifts. These variants also show dynamic frequency changes across time points—for example, CLIPA9 Phe233Tyr rises from 75.6% in infected mosquitoes at Day 1 to 58.3% at Day 17, while remaining below 50% in uninfected pools; LRIM17 Pro401Ala remains near fixation in infected pools (>97%) but increases in uninfected pools over time. Notably, CLIPB3 is absent from the heatmap because its differentiation was driven by synonymous SNPs rather than amino acid changesThese frequency patterns and statistical associations support the hypothesis that vector competence is shaped by a dynamic and multilayered immune adaptation to infection.

**Fig 4.**
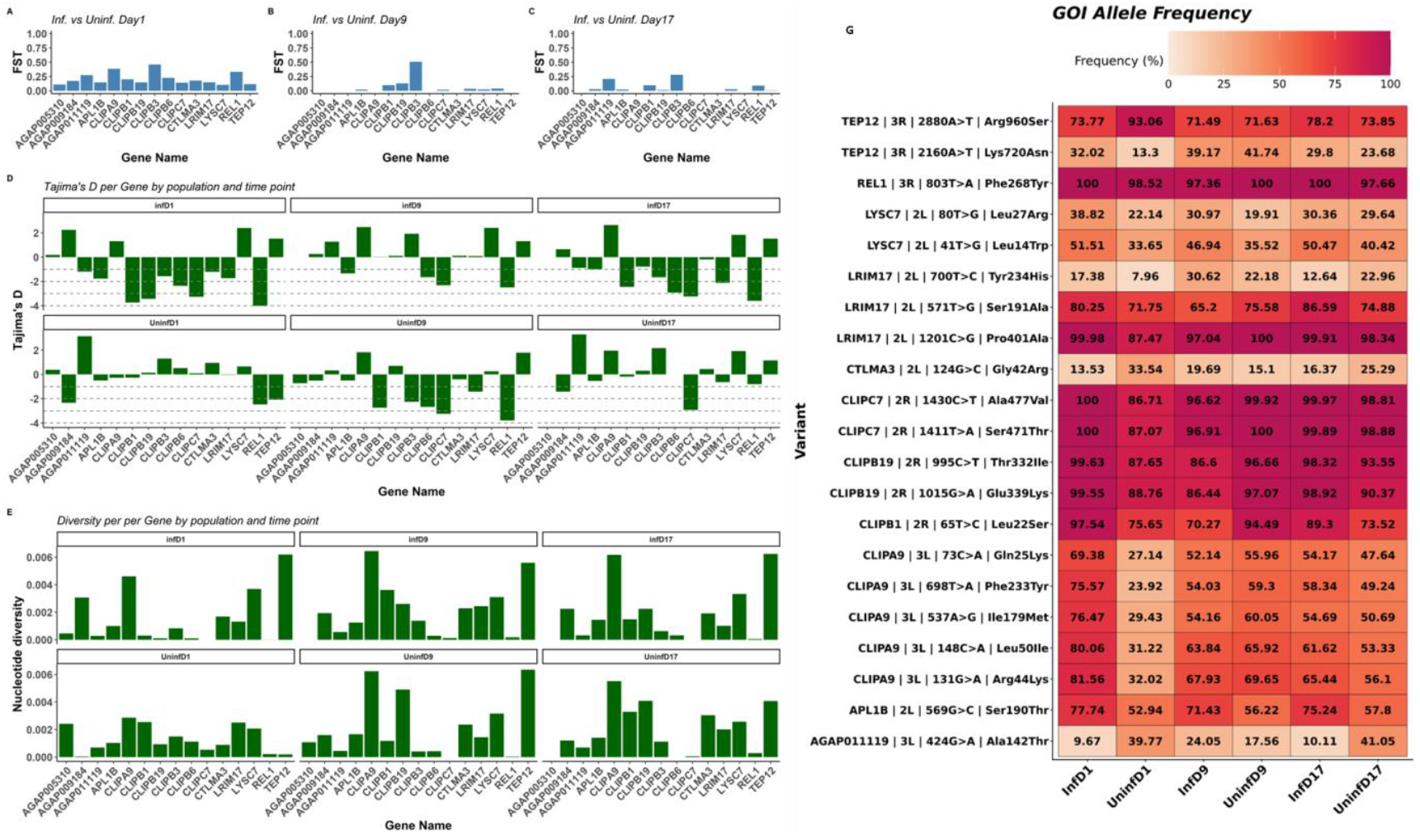
Genetic differentiation and diversity metrics for candidate genes in infected and uninfected An. gambiae mosquitoes across time points. Panels (A–C) show gene-specific FST values comparing infected (Inf) versus uninfected (Uninf) mosquitoes at Day 1 (A), Day 9 (B), and Day 17 (C). Only genes with FST ≥ 0.1 between Inf and Uninf at Day 1 are displayed, as this threshold was used to select candidates showing substantial genetic differentiation. Panel (D) presents Tajima’s D per gene across infection status and time points, and panel (E) shows nucleotide diversity (π) per gene across the same conditions. Panel (F) is a heatmap illustrating allele frequencies of nonsynonymous SNPs detected in these candidate genes. Frequencies are color-coded from light orange (low) to dark red (high). Gene names, chromosomal positions, and amino acid substitutions are listed on the left.

## Discussion

The immune responses of *An. gambiae* against infection by *P. ovale* has not been characterised to date. In this study, we provide the first transcriptome analysis of *An. gambiae-kisumu* infected by *P. ovale* at the different development stages (ookinete, oocyst and sporozoite). Our analysis identified several genes that are either well-established or strongly implicated in mosquito immune responses and parasite transmission-blocking mechanisms. We also identified several potential novel candidates that could influence vector competence and parasite development.

### Temporal Gene expression patterns during *P. ovale* infection

The pairwise comparison within the different time-points revealed that *Anopheles gambiae_kisumu* transcriptional responses to *Plasmodium ovale* infection were more active during the ookinete formation (24h post infection) than the other stages. This pattern is similar to the general activation of strong immune response in mosquitoes when *Plasmodium* crosses the peritrophic matrix and invades the gut epithelium (36, 37).

Among differentially expressed genes (DEG) in *P. ovale*-infected mosquitoes, CTL3, CLIPB12 and LRIM10 were among the top immune overexpressed genes. CLIPB12, from the clip domain serine proteases family, exhibited significant overexpression (FC 49.9) during *P. ovale* infection. This gene has not been fully characterized. This high level of upregulation appears to be specific to the response against *P*.*ovale*, particularly at the ookinete development. This is consistent with the established functions of other CLIP protease family members such as CLIPB8, CLIPB9, CLIPB10, CLIPB14 and CLIPB15 which have been shown to play a role in extracellular proteolytic cascades that mediate the killing and lysis of *P. berghei* ookinetes and Gram negative bacteria in Anopheles mosquitoes (38, 39). Interestingly, the CLIPB15 was found significantly downregulated at 9 dpi. LRIM10 although its role has not been previously characterized, other LRIM such as LRIM1 and LRIM9 have been well studied and have been functionally linked to innate immunity and killing of Plasmodium ookinetes in the midgut epithelium (40). The observed high expression level of LRIM10 with 11.2 fold change could be due to the activation of immune mechanisms to fight against *P. ovale*.

Thirteen genes with functions predicted to be involved in calcium ion binding showed higher expression levels in infected mosquitoes 24 hours post infection. Among them Troponin C (AGAP006181) was the most significantly expressed with up to 85 fold change. Troponin C is a calcium-binding protein that plays an important role in the regulation of muscle contraction by binding calcium ions (Ca^2+^) (41, 42). Moreover, Calcium is known to be essential for ookinete motility (43). These genes alterations may supply parasites with Ca2+ which may facilitate midgut invasion. High expression level found in infected mosquitoes only at ookinete stage needs to be further investigated to establish if mosquito Ca^2+^homeostasis can directly influence *Plasmodium* infectivity and whether *Plasmodium* parasites directly alter calcium concentration in midgut epithelial cells.

The low transcriptomic response to *P. ovale* infection observed at 9 dpi compared to other time-points could indicate a relatively lesser immune response at the late oocyst production stage. In fact, the oocyst maturation, between one and two weeks depending on the *Plasmodium* species correspond to the period where oocysts grow in size and undergoes sporogonic division and end with sporozoite formation. In our study, two genes (heme peroxidase 3 (HPX3) and vitellogenin (Vg)) emerge as potential modulators of the parasite-vector interactions at 9 dpi. Previous studies have shown that heme peroxidase, HPX 2 and NADPH oxidase 5 (NOX5) in *An. gambiae* contribute to *P. berghei* clearance through nitration of epithelial cells (44). The upregulation of HPX3 observed in our data suggests a similar immune effector role, possibly contributing to midgut epithelial defense against parasite invasion. In contrast, vitellogenin (Vg) showed sustained overexpression at all time points. Vg is a major yolk protein precursor produced by the fat body during vitellogenesis (egg production) and has been showed to facilitate *Plasmodium* survival by helping the parasite to evade the mosquito immune defense (45).

### Identification of potential candidate genes blocking transmission to validate through GO enrichment analysis

GO enrichment analysis of top differentially expressed genes in response to *P. ovale* infection revealed the involvement of immune pathways and effector functions relevant to blocking malaria parasite development and transmission. Several of the most prominent differentially expressed genes including LRIM1, TEP1, CLIPB12, CLIPC9, LRIM10, and PPO6 share GO terms associated with immune response (GO:0006955), proteolysis (GO:0006508), and extracellular region (GO:0005576). These genes play an important role in melanisation and complement-like pathways, which encapsulate and kill invading parasites, thereby serving as a natural transmission-blocking mechanism or for mosquitoes or for transgenic flies (36, 38).

It has been shown that LRIM1 and TEP1 are part of the mosquito complement-like pathway, facilitating the recognition for effective lysis or melanization of *Plasmodium* ookinetes, significantly reducing parasite survival in the midgut (30, 46). Similarly, CLIPC9, serine proteases are involved in proteolytic cascades that activate melanization, a process that encapsulates and kills invading parasites (18, 46). PPO6 (prophenoloxidase 6) is a key enzyme in the melanization pathway, further contributing to parasite clearance. The upregulation of LRIM10 and CLIPB12, which share GO terms for immune response and protein binding with LRIM1, CLIPC9 and TEP1 suggests a strong involvement of these genes in immune responses against *Plasmodium*.

The APN gene family (APN1 and its paralogs (APN2, APN3, APN4, APN5)), all sharing GO terms for proteolysis (GO:0006508) and metallopeptidase activity (GO:0008237), is particularly noteworthy. APN1, have been already established as promise transmission-blocking vaccine (TBV) target due to its role in parasite invasion of the mosquito midgut (37, 47, 48). The upregulation of its paralogs suggests a coordinated function in midgut physiology and parasite defense.

Furthermore, genes involved in detoxification and oxidative stress response, such as the cytochrome P450, CYP9K1 and the glutathione S-transferase, GSTD3, were also consistently upregulated at different time-points (9 and 17 dpi), sharing the GO term for oxidoreductase activity (GO:0016491). Recent studies have shown the implication of metabolic resistance genes including P450s (CYPs and GSTs, not only involved in detoxification of insecticides but are also associated with higher Plasmodium infection rates (49-51). It has been shown that the L119F-GSTe2 mutation in *Anopheles funestus* is linked to increased sporozoite infection and higher malaria transmission intensity (52). Similarly, overexpression of GSTs and P450s has been correlated with increased susceptibility to *Plasmodium* in several vector populations (51, 53). Changes in expression of detoxification genes during parasite infection in vectors suggest a potential role, although more study is needed to confirm their utility as transmission-blocking targets.

ABCC9, an ATP-binding cassette transporter, was upregulated and shared GO terms for membrane localization (GO:0016020) and transporter activity (GO:0005215) with APN1, suggesting a role in modulating the midgut environment or in the transport of immune effectors. Another ABCG2, mainly involve in *Plasmodium* development has been shown essential for female gametocyte and zygote formation (54).

### Genes involved in mosquito immune responses exhibit strong genetic differentiation and signatures of selection early after infection

Our findings highlight the importance of genetic variation related to the immune system in determining mosquito susceptibility to *P. ovale*. The non-synonymous mutations identified in gene such as *APL1C, LRIM15, LRIM17*, and *CTLMA2* suggest that even minor amino acid substitution could have a significant impact on protein structure, signaling dynamics or interaction with parasites-derived molecules. Variants such as Thr567Ala and Glu330Val, which alter the polarity or charge, may affect folding or stability of immune complexes, thereby modulating the efficiency with which parasites are recognised and cleared. Similarly, amino acid change to Ala116Thr in *CTLMA2* and Pro401Ala in *LRIM17* demonstrated how structural changes in pattern recognition or effector protein can influence immune responsiveness. The reccurence and persistence of certain mutations, particularly within APL1C, across multiple time points to their involvement in both the early and sustained phase of mosquito immune response. These observations are consistent with previous studies showing that genetic variation shapes mosquito immune pathways, particularly, those envolving complement-like and leucine-rich repeat proteins, and that these pathways play a critical role in determining vector competence (18, 55). Furthermore, as observed in human studies where cytokine gene polymorphisms influence malaria outcomes, the presence these polymorphisms indicates that similar principe of immune modulation may exist across host and vector species (56). Taken together, these findings suggest that vector competence is not governed by a single genetic determinant, but rather by a coordinated network of immune-modulating variants whose effects evolve alongside vhanges in gene expression throughout the course of infection.

## Conclusion

This study provided an overview of the mosquito immune responses to the infection by *P. ovale* identidying key genes controlling this process at different time-points. The overexpression of many of these genes during *P. falciparum* infection against *An. gambiae* highlights conserved immune pathways that could be exploited to reduce malaria transmission. Several of the newly identified candidates are promising targets for transmission-blocking strategies and functional validation.

## Acknowledgements

We thank the local residents of Elendé village for their participation in this study.

## Author contributions

DNN, CSW, SB, AAA, designed the study. DNN, FNN, CN performed experimental infection. DNN performed lab analyses and RNAseq. DNN, CSW, AT, APY analysed data and prepared Fig, interpret the results and wrote the manuscript. DNN wrote the first draft. DNN, CSW, AT, APY, FKN, CN, STBS, SB, AAA, FNN read commented and approved the manuscript.

## Supporting information

**S1 Fig. *Plasmodium ovale* oocyst observed at day 12 post infection**.

**S2 Fig Distribution of SnpEff-annoted Reads by Functional Class and Genomic Region**. Panel A illustrates the proportion of reads categorized by functional class_Nonsense, Misense and Silent_ across various regions, indicating a predominance of Silent mutations. Panel B displays the distribution of reads across different genomic regions, highlighting the most common sites such as Exons, UTRs and splice sites, with a ntable enrichment of certain annotations like UTRs and splice sites in specific regions.

**S1 Table. List of Primer**.

**S2 Table. Summary of Sequencing Quality Metrics Across All Samples**.

**S3 Table. Upregulated genes InfD1.vs.UnInfD1**. InfD1: mosquito infected at Day1; UnInfD1: Uninfected Day1

**S4 Table. Genes expressed at different time points**

**S5 Table. Candidate mosquito genes sharing enriched Gene Ontology terms**

